# Triazavirine supramolecular complexes as modifiers of the peptide oligomeric structure

**DOI:** 10.1101/150664

**Authors:** Alexey V. Shvetsov, Yana A. Zabrodskaya, Peter A. Nekrasov, Vladimir V. Egorov

## Abstract

In this study we present molecular dynamics simulations of the antiviral drug triazavirine, that affects formation of amyloid-like fibrils of the model peptide (SI). According to our simulations, triazavirine is able to form linear supramolecular structures which can act as shields and prevent interactions between SI monomers. This model, as validated by simulations, provides an adequate explanation of triazavirine’s mechanism of action as it pertains to SI peptide fibril formation.

## Introduction

Conformational changes play a significant role both in normal protein functioning and in disease. Amyloid-like fibril formation is the process that underlies the pathogenesis of conformational diseases. Such fibrils are resistant to proteolytic enzymes, as well as to chemical and physical influences [16]. Amyloid-like oligomers can also act as signal transducers in health and in disease [19]. Agents that affect protein oligomer formation can be used as fibrillogenesis inhibitors, or as signal transduction regulators. In our previous studies [6], we demonstrated the effect of triazavirine (a guanine nucleotide analog [11], TZV, Figure 1) on GDIRIDIRIDIRG peptide (self-complementary ionic, SI) fibril dissociation. Transmission electronic microscopy and small-angle neutron scattering were effectively used to study the interaction.

**Figure 1:**
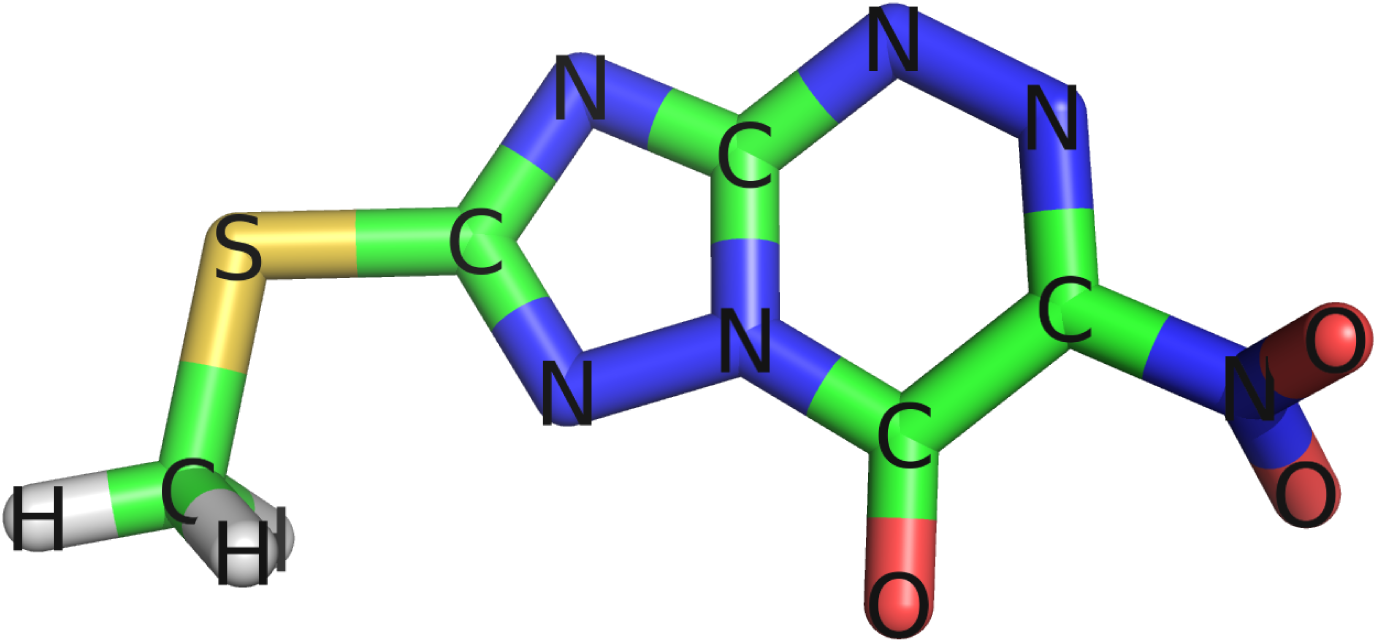
Structure of triazavirine (a guanine nucleotide analog [11], TZV)

The SI peptide is comprised of an ionic, self-complementary motif (iSCM). This motif was first identified in small heat shock proteins, crystalline oligomeric complexes [7], and in peptides with labile secondary structures [2]. The interaction between these polypeptides occurs at a site where antiparallel-oriented iSCMs monomers (on opposing polypeptides) interact.

Electrostatic attraction between monomers, and zipper-like hydrophobic interactions between oligomers, are the two major driving forces allowing iSCM to successfully complete the self-organization process [4], [10]. In previous studies [6], we hypothesized that triazavirine, due to its status as a guanosine analogue, may have a tendency to interact with arginine [12]. Triazavirine’s interaction with iSCM most likely involves a shielding of arginine’s positive charge by the drug, and a modification of the motif’s overall charge. The modified local charge environment shifts the balance towards protein aggregate dissociation and towards monomer formation (via repulsion). Molecules with similar anti-fibrillogenic mechanisms of action have been described previously [15, 14]. In this study, we examined triazavirine with molecular dynamics simulations in terms of its action on SI peptide fibrils.

## Materials and Methods

### Modelling of the TZV system

TZV modelling was accomplished using the Avogadro[8] software package. The topology of TZV for GROMACS was generated using ACPYPE[17] and antechamber from Ambertools[18]. Two systems with 32 and 320 TZV molecules in 1000nm^3^ boxes (10x10x10 nm) were created using gmx insert-mol.

### Modelling of the SI peptide system

Initial conformations of GDIRIDIRIDIRG peptide were constructed as unfolded states using Pymol[5]. After that, 16 GDIRIDIRIDIRG peptide chains were inserted into 1000nm^3^ box (10x10x10 nm) in random orientations using gmx insert-mol. In models which included TZV in addition to 16 GDIRIDIRIDIRG molecules, 320 TZV molecules were inserted with random orientations using gmx insert-mol.

### Molecular dynamics

All MD simulations were performed in the GROMACS [1, 3] software package using amber99sb-ildn[13] force field for peptides, and tip3p[9] model for explicit water. Simulated systems were around 140k atoms in size including explicit solvent shell and 50mM NaCl to neutralize charges on peptides (and TZV, if present in the model). Solvent shell thickness was at least 2.5 nm. The systems were equilibrated using a two step protocol. During the first step, the system was equilibrated for 5 ns with all heavy, non-solvent atoms restrained to their initial positions using NPT ensemble. During the second step, the system was equilibrated without restraines for 10 ns (starting from the last frame of the previous step). After two equilibration stages, a 500 ns trajectory was simulated with a time step of 2 fs. A Neighbour search was performed every 50 steps. The PME algorithm was used for modeling electrostatic and Van der Waals interactions with a cut-off of 1.2 nm. Temperature coupling was done with the Nose-Hoover algorithm at 310K. Pressure coupling was done with the Parrinello-Rahman algorithm for 1 bar.

### Quasi-elastic light scattering

Quasi-elastic light scattering (QLS) measurements were performed using QLS-spectrometer LCS-03 (Intox, Russia). Spectra analysis was performed with the spectrometer software using an “rod-like” model of scattering objects.

## Results and discussion

Molecular dynamics simulations showed that the SI peptide in solution is prone to the formation of aggregates. Figure 2a shows the final condition of 16 SI peptide molecules after 500 ns of simulation.

**Figure 2:**
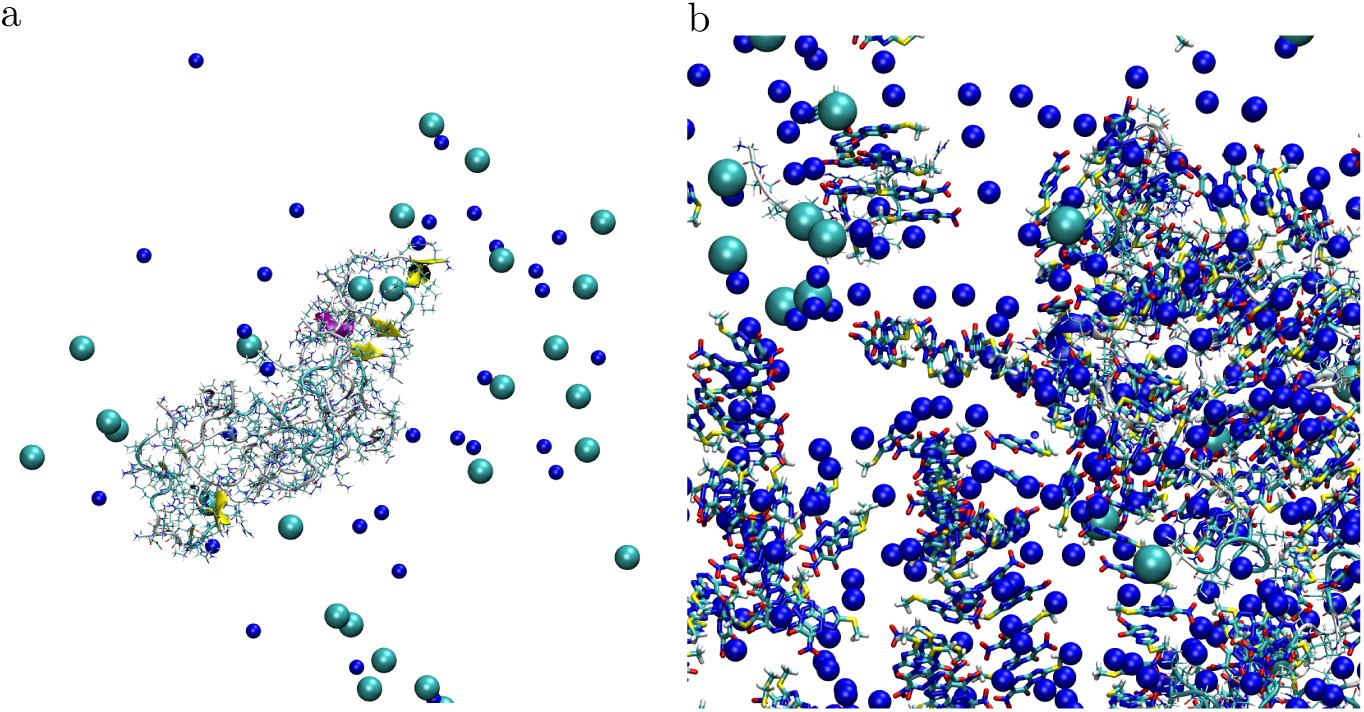
Final state of a 500 ns molecular dynamics simulation of 16 SI peptide molecules in solution (a) and 16 SI peptide molecules in solution with 20-fold molar excess of triazavirine (b)

Interestingly, the molecular dynamics simulations of triazavirin in solution also showed that it has a tendency to form linear supramolecular complexes at concentrations above 10 mg/ml. Figure 3 presents the results of a simulation of an ensemble of 32 triazavirin molecules (equivalent to 11 mg/ml) and 320 triazavirin molecules (equivalent to 110 mg/ml) in a 1000 nm^3^ cell.

**Figure 3:**
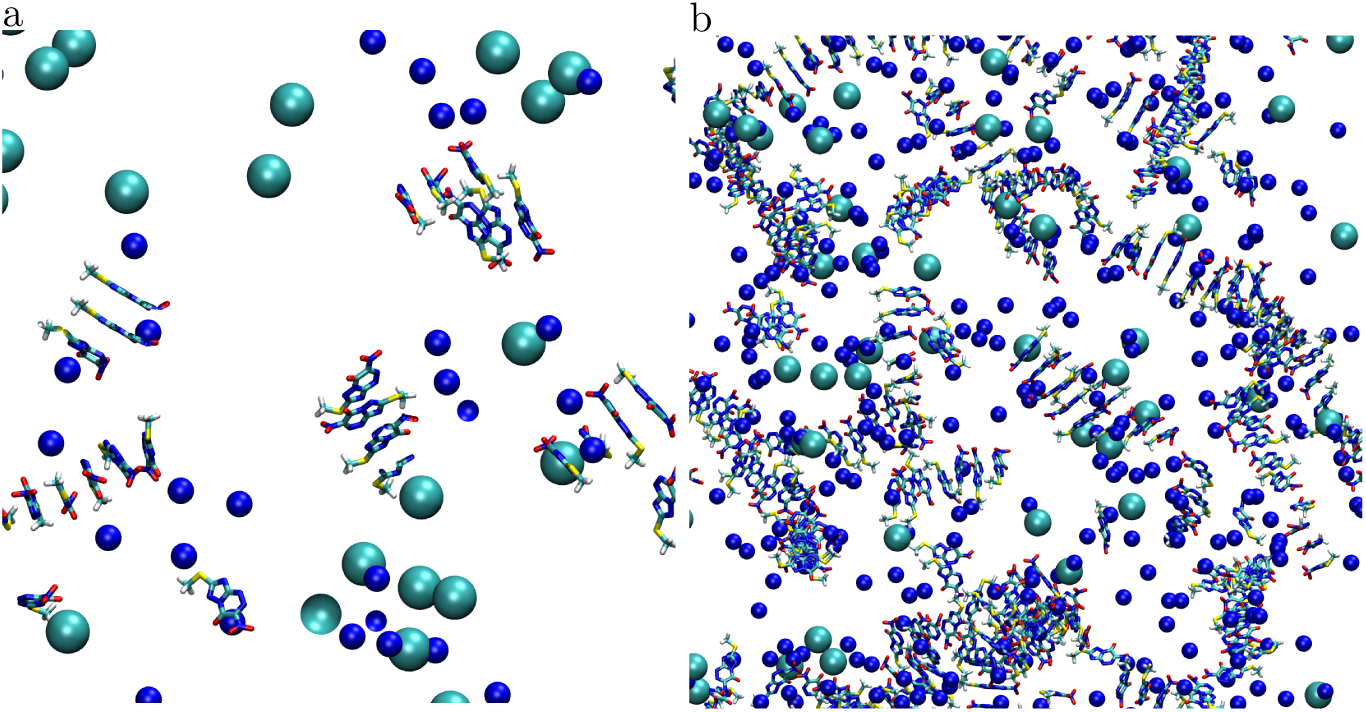
The final state of a 500 ns molecular dynamics simulation of 32 (a) and 320 (b) triazavirin molecules in solution. There is a tendency for the formation of linear structures.

It should be noted that during in-vitro measurements using QLS in triazavirin solutions (after filtering), we also observed the presence of light-scattering structures at concentrations above 2 mg/ml. Figure 4 shows the results of QLS of triazavirin solutions at concentrations of 2 mg/ml and 20 mg/ml.

**Figure 4:**
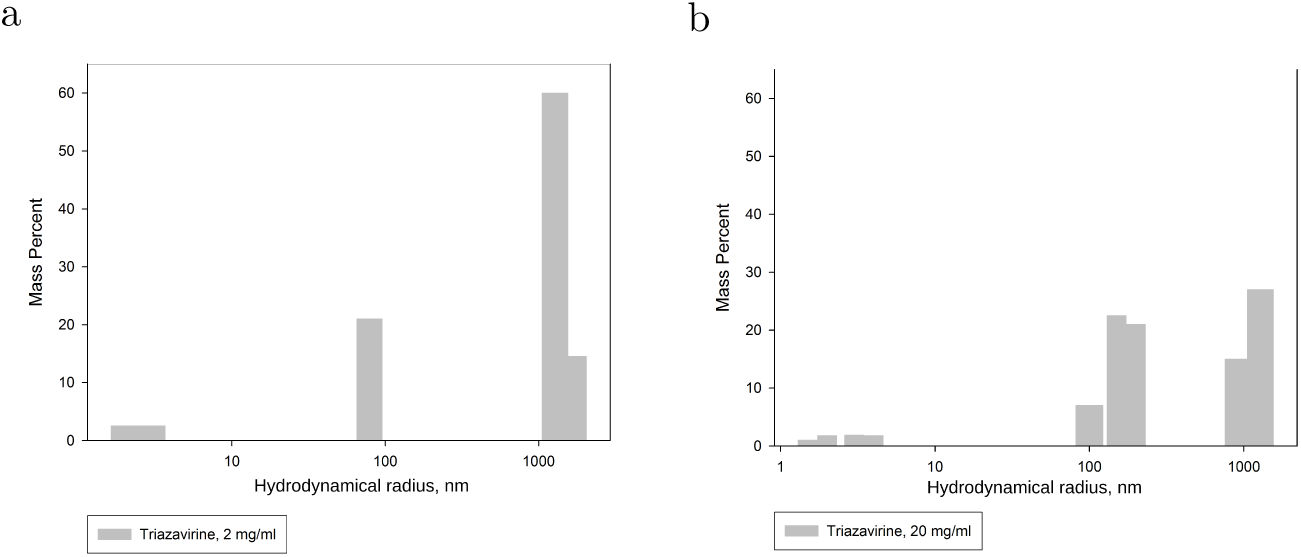
Results of QLS measurements of triazavirin solutions at 2 mg/ml (a) and 20 mg/ml (b) concentrations.

In QLS experiments, the formation of sub-microscopic particles in triazavirin solutions was observed. The triazavirin critical concentration in solution (below which particle formation was not observed) was 2 mg/ml. We performed measurements immediately after dilution, after 30 minutes, and after 60 minutes. Particle formation dynamics did not depend on the tested concentrations. Immediately after preparation, a fraction of particles with a hydrodynamic radius of about 100 – 200 nm was observed. This can be explained as dissolution of the drug, in its crystalline particle form, into aqueous solution. Then, at the next measurement time point (30 minutes), the fraction was split into 2 sub-fractions. Further changes in fraction size distributions (after 60 minutes) were not observed. The hydrodynamic radius of the particles at concentrations of 2 mg/ml were 75-90 and 1000-1500 nm for concentration of 2 mg/ml, and 90-250 and 600-1500 for concentration of 20 mg/ml.

Thus, the QLS experiments have shown triazavirin’s ability to form supramolecular complexes. In order to study the influence of such complexes on the formation of fibrils in solution, molecular dynamics simulations of the SI peptide in the presence of a 10-fold molar excess of triazavirin were performed.

Figure 2b shows the final state of 16 SI peptide molecules in the presence of 320 triazavirin molecules after 500 ns simulation.

A comparison of pairwise distances between the peptide molecules’ center of mass for SI with and without TZV reveals an increase in the average value of pairwise distances. In Figure 5 there is a colored diagram of such a comparision.

**Figure 5:**
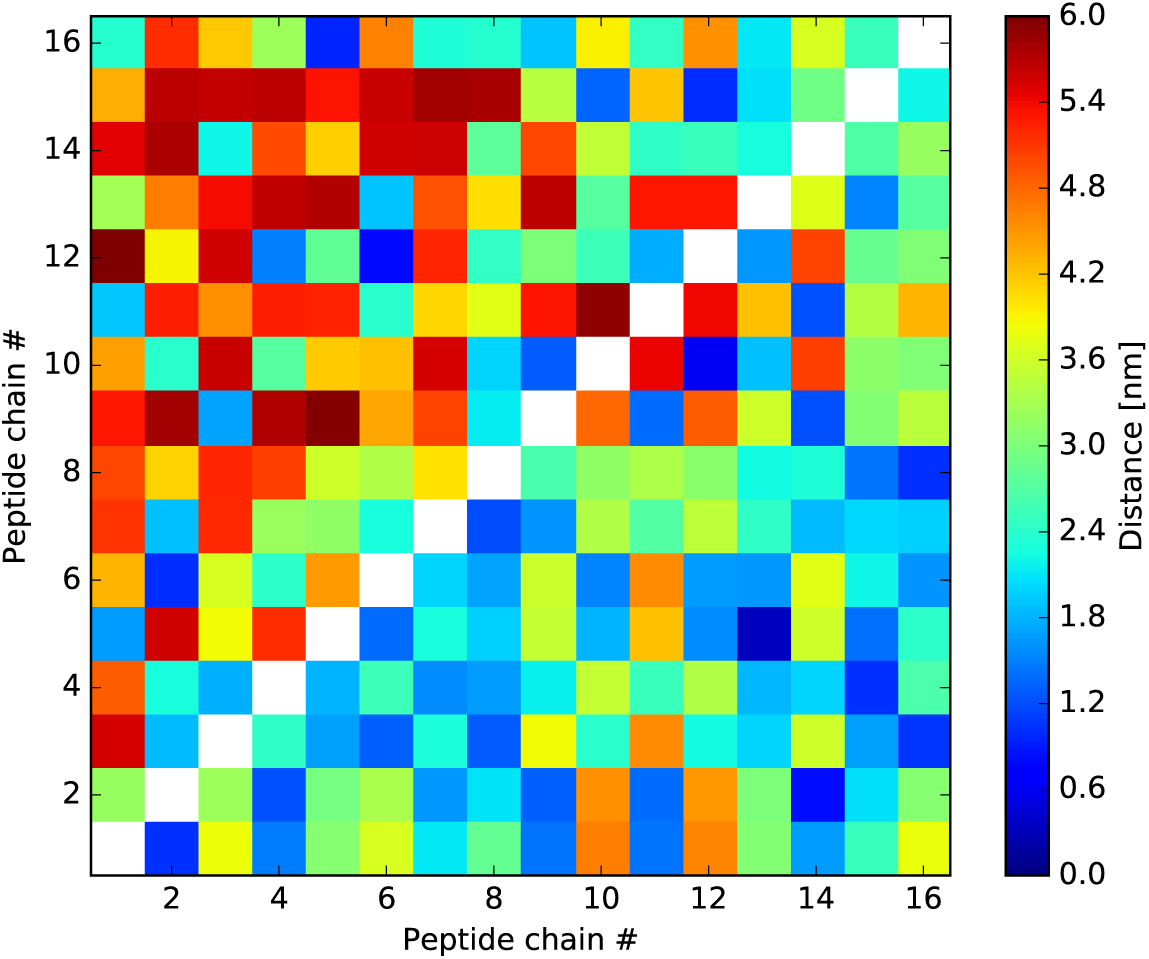
Diagram of pairwise distances between SI peptide molecules in the process of simulation in the presence (above diagonal) or absence (below diagonal) of 20-fold triazavirine excess

Figure 6 shows that rod-like TZV supramolecular complexes act as shields and prevent peptide-peptide interaction. In Figure 7 there is a histogram of mean center of mass pairwise distances between SI peptide and TZV molecules. It showed that some of TZV molecules have stable positions near SI peptide monomers during the last 250 ns of simulation.

**Figure 6:**
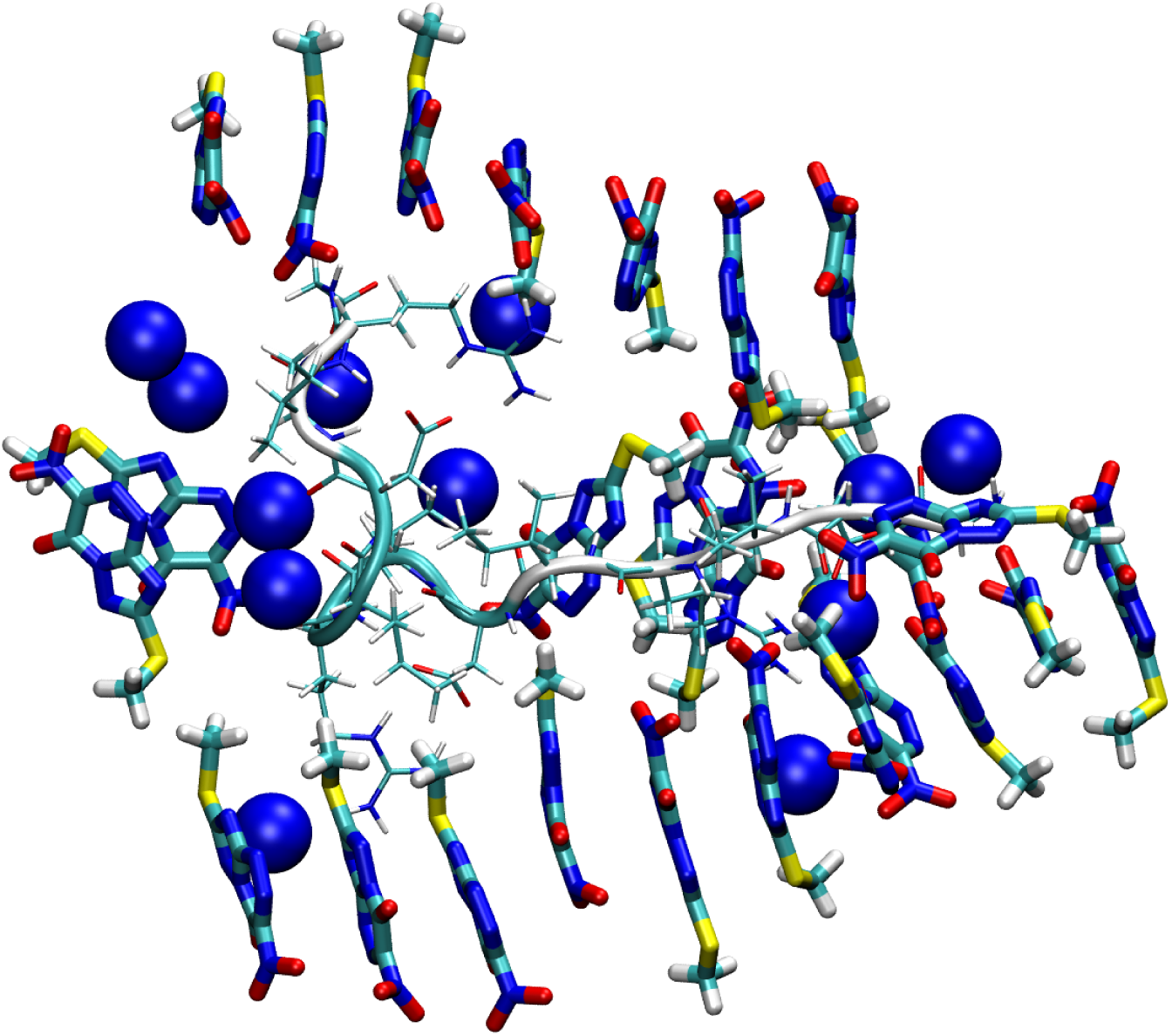
Interactions between SI and TZV supra-molecular complexes

**Figure 7:**
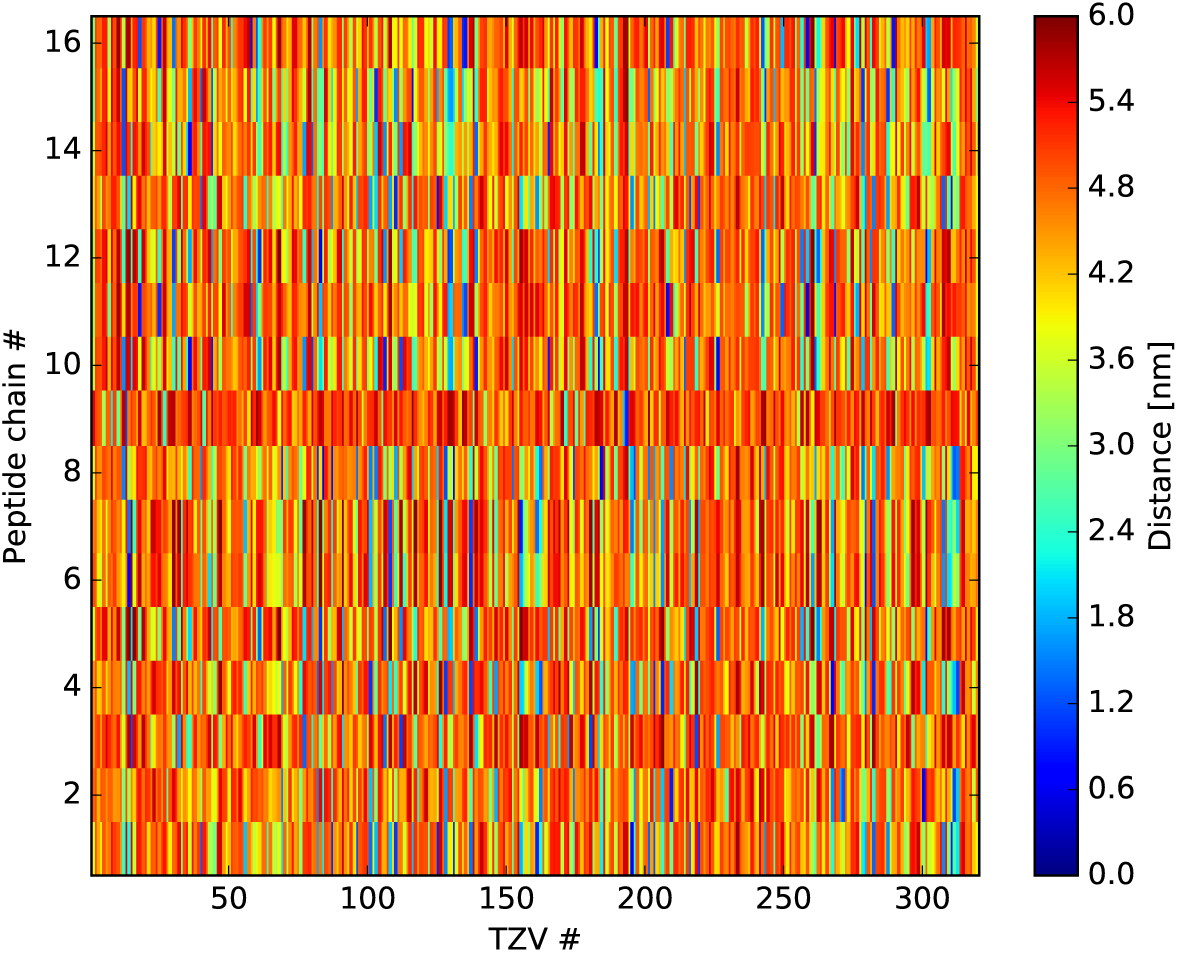
Diagram of mean pairwise distances between SI peptide molecules and TZV molecules during last 250 ns of simulation

In silico simulation shows that triazavirine, at 10 mg/ml and higher concentrations, can form supra-molecular linear structures. The presence of such structures in solution was detected by QLS, and the results are consistent with our previous small-angle neutron scattering data. According to our simulations, these linear triazavirine supramolecular structures act as shields and prevent interactions between SI monomers. Our model, as validated by simulations, provides an adequate explanation of triazavirine’s mechanism of action as it pertains to SI peptide fibril formation.

## Conclusion

Thus, the molecular dynamics simulation shows that, apparently, in addition to interactions between triazavirin molecules and side chains of arginines (described in [6, 16]), interactions between peptide molecules and triazavirin supramolecular complexes can occur.

## Acknowledgments

The results of the work were obtained using computational resources of Department of Information and Computational Technologies of the St. Petersburg State Polytechnic University (<u>http://www.spbstu.ru</u>). We thanks Mr. Edward Ramsay for the help in writing the manuscript and Dr. D. V. Lebedev (DMRB PNPI NRC KI) for the fruitful discussion of the experiments. This study was supported by the Russian Foundation for Basic Research, project no 14-24-01103 ofi_m.

